# Functional Social Structure in Baboons: Modelling Interactions Between Social and Environmental Structure in Group-Level Foraging

**DOI:** 10.1101/270041

**Authors:** Tyler R. Bonnell, S. Peter Henzi, Louise Barrett

## Abstract

In mobile social groups, cohesion is thought to be driven by patterns of attraction at both the individual and group level. In long-lived species with high group stability and repeated interactions, such as baboons, individual-to-individual attractions have the potential to play a large role in group cohesion and overall movement patterns. In previous work, we used GPS mapping of a group of baboons in De Hoop, Western Cape, South Africa, to demonstrate the influence of such attractions on movement patterns. We also demonstrated the existence of emergent group-level structures, which arose as a consequence of individual social influence. Specifically, we found a core-periphery structure, in which a subset of influential animals exerted an influence on each other and those animals in the periphery, while those in the periphery were influenced by the core but did not exert any influence over others. Here, we use agent-based modelling of baboon groups to investigate whether this group-level structure has any functional consequences for foraging behaviour. By varying individual level attractions, we produced baboon groups that contained influence structures that varied from more to less centralized. Our results suggest that varying centrality affects both the ability of the group to detect resource structure in the environment, as well as the ability of the group to exploit these resources. Our models predict that foraging groups with more centralized social structures will show a reduction in detection and an increase in exploitation of resources in their environment, and will produce more extreme foraging outcomes. More generally, our results highlight the link between social and environmental structure on functional outcomes for mobile social groups of animals.

## Introduction

Among the primates, group living is thought to have evolved as a means to reduce predation risk, and competition within and between groups is thought to influence a group’s social structure (Boinski and Garber, 2000; Van Schaik, 1983, 1989). Baboons have long been used to test such socioecological theories, because they are one of the best studied of all primate taxa and occupy a wide range of ecologies (Henzi and Barrett, 2003; Henzi and Barrett, 2005). Efforts have also been made to study the internal structure of baboon groups at a more proximate level. In a now classic paper, Stuart Altmann (1974) made a series of predictions regarding the manner in which resource distribution and competition would structure the geometry of baboon groups. Thanks to advances in technology that allows individual positions to be mapped, some of Altmann’s insights have now been tested, and his predictions have been shown to apply in at least one baboon population (Dostie et al., 2016). High resolution sampling of behaviour in other baboon populations has also shown how resource distribution and social interactions between animals combine to determine the geometric structure of groups (Farine et al., 2016).

Using similar methods, it has also become possible to capture the “social influence” structure of groups from empirical data (Bonnell et al., 2017; Eriksson et al., 2010; Katz et al., 2011; Lukeman et al., 2010; Mann, 2011). That is, how patterns of attraction and repulsion between individuals give rise to the internal structure of groups. In our own work on baboons, we have considered how group-level structures can arise from the combination of individual influence patterns (Bonnell et al., 2017). Specifically, Bonnell et al. (2017) found evidence for a core/periphery structure at the group level, where a core of more dominant, inter-dependent individuals exerted a unidirectional influence on the movements of other, peripheral animals.

An obvious question that arises from such findings is whether any functional benefits accrue from particular influence structures. Research to date has shown that local influence between neighboring individuals can propagate information through collectives faster than any individual can travel (Sumpter et al., 2008). Similarly, there is evidence that such local interactions allow a few knowledgeable individuals to guide the decisions of a large number of naive individuals (Couzin et al., 2005). In these cases, the effects of influence structures are dominated by spatially-neighbouring individuals, where all individuals are treated as homogenous and have equal influence. In cases like this, group size alone may prove to be an advantage in collective decision making (referred to as the “wisdom of the crowd”: (Galton, 1907). This occurs through the averaging of individual decisions, resulting in group decisions closer to optimal than any one individual. When there is internal structure to a group, however, the specific network of connections between individuals can influence group decision making (Krause et al., 2010; Rosenthal et al., 2015). It is also important to recognize that such internal structure can give rise to emergent patterns that do not necessarily confer an advantage. That is, emergent patterns can often result simply from the existence of non-linear interactions (Bradbury and Vehrencamp, 2014). Consequently, it is important to consider what, if any, advantage a particular pattern might convey, and in what contexts (Parrish and Edelstein-Keshet, 1999).

Here, we develop testable predictions about the functional role of influence structures within mobile simulated baboon troops engaged in foraging tasks that can be applied to real-world situations. This will enable more precise predictions regarding the influence of habitat structure and composition on group shape and structure across baboon populations, as well as contributing more generally to work in movement ecology and collective behaviour.

To explore the functional consequences of variation in a core-periphery structure we use agent-based modelling. Specifically, we investigate how characteristics of the resource landscape interact with internal group structure to promote or impede the ability of groups to locate resource-rich areas, and subsequently take advantage of them. We expected to find that less centralized social structures (i.e., those with a larger core of influential animals) will result in (i) the group as a whole being better able to identify high value resource structures on the landscape, and (ii) will result in less within-group variance in foraging efficiency. In more centralized groups (i.e., those with a smaller core), we predicted the opposite trends.

To achieve this, we quantify the foraging efficiency of simulated groups by performing virtual foraging trials. In these trials, we alter the social influence structure of the group, the size of the group, and the structure present in the resource landscape. We define influence structures within these simulated groups using a core-periphery approach, where a core is defined as a set of inter-dependent individuals, and peripheral individuals are those that are influenced by the core but not each other (Fig. 1). We varied influence structures by altering the size of the core, generating influence structures ranging from a single leader (e.g., one individual is the core) to a homogenous influence structure (i.e., all individuals form part of the core) (Fig. 1). We further varied group size to alter the magnitude of scramble competition. Finally, we altered the resource landscapes in which our foraging experiments were run, creating a context where resources were distributed randomly and homogenously, versus a context in which a single high-density resource path was present and one in which several high-density paths were present. We used a single high-density path in order to provide a clear optimum for foraging so that we could quantify the relative effects of social influence structure and group size on the ability to exploit environmental structure.

**Figure 1:**
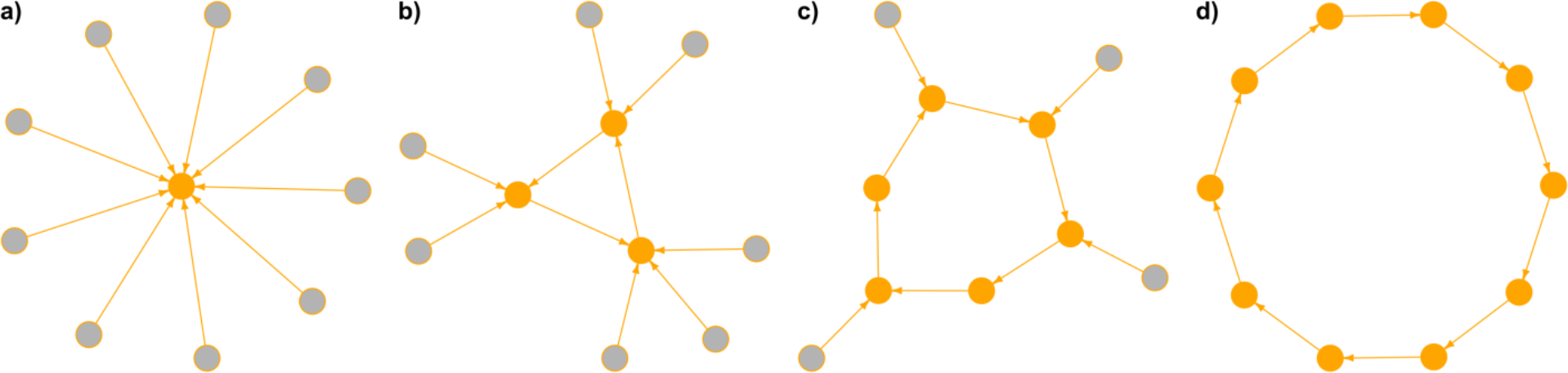
Group level influence structures in four groups of 10 individuals: a) one individual at the center (C_per_ = 0.1), b) three individuals form a core (C_per_ = 0.3), c) six individuals form a core (C_per_ = 0.6), and finally d) all individuals are inter-dependent (C_per_ = 1.0).

## Methods

### Movement model

The movement model used is based on correlated random walk models (Van Moorter et al., 2009). In our model, animals are simply biased towards visible sites that are close and have high resources. To calculate the resulting influence of food patches on a simulated animal, we weight each patch within a visual radius (R_vis_=50m) based on the distance from the focal animal and the amount of food at that patch, 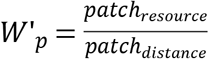. Where patch resources vary from 0-1. We then standardize the patch weights to sum to one, 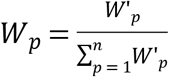, and calculate the average food vector based on these weights 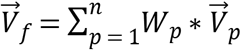.

Along with this motion bias towards resources, we add a social attraction force into the model by adjusting motion based on attraction to a particular group member. We use a linear function describing an increasing attraction towards a group member beyond an attraction radius (d_a_=10m) (Couzin et al., 2002; Warburton and Lazarus, 1991):

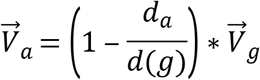

The attraction vector 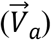 of the focal animal describes the attraction to one other individual. The combined result of these forces are thus:

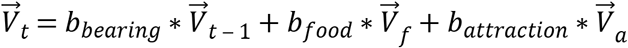

Where 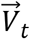 is the resulting motion vector at time t, 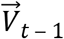 is the previous motion vector, 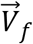 is the vector towards food patches, and 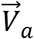 is the attraction vector. The parameters *b*_*bearing*_, *b*_*food*_, and *b*_*attraction*_ represent the relative influence of each force acting on the simulated animal. We set *b*_*bearing*_ and *b*_*food*_ to a value of 1, and *b*_*attraction*_ to a value of 2. This produces a set of conditions where social forces predominate over food or movement persistence, and where movement persistence might be expected to be relatively similar to food bias, i.e., under conditions where food is of low value and widely distributed.

To account for variable uncertainty in motion due to conflicting forces, the final resulting motion vector is sampled from a wrapped normal distribution (Von Mises) with 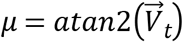, and 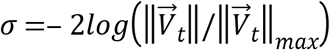. Where 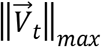 is simply the maximum length possible of the resulting influence vectors (e.g., when they all point in one direction). This results in very little uncertainty around 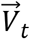 when all the influencing factors are operating in the same direction and increased uncertainty in motion when they are all conflicting (Van Moorter et al., 2009).

### Social influence structures

Each group added to a foraging trial is initialized with a fixed influence structure, where each individual is assigned one other group member to “follow.” These influence structures are defined by assigning individuals to either core or periphery status. Each group is assigned a group size (G_size_) and a percentage of individuals in the core (C_per_). By varying these parameters, we can create influence structures that are more or less despotic or democratic (Fig. 1). The larger the core size in the group, the more foraging decisions represent the outcome of many interdependent movements. Conversely, the smaller the core group, the more the group foraging decisions are “despotically” driven by one individual’s movements.

### Simulated foraging trails

First, we investigated the influence of varying core and group size on foraging behaviour in a uniform versus heterogeneous landscape. We simulated a base landscape (2000m × 2000m) with randomly distributed food patches (0.01 patches/m^2^), assuming a homogenous resource landscape with opportunistic and quickly depleted patches. We compared to this to a second landscape that contained a high food density path with twice the number of randomly distributed patches (0.002 patches/m^2^). This path was non-linear and follows a parabolic curve, starting at the bottom-left corner of the landscape. We used a uniform random distribution to generate 1000 groups with group size varying between 5 and 100 agents, and the proportion of group members constituting the core varying from 0 to 1. Each group was then run on both the path and non-path landscapes.

Each simulation starts a group at the bottom-middle of the landscape and allows the group to forage for 2 hours (7200 time steps). The 2-hour limit marks the approximate time that a large group traveling along the high density path would take to reach the top of the simulated landscape, thus depleting the high resource path and rendering the resource landscape equivalent to the non-path environment. By constraining the time to 2 hours, we focus on the time period where the path and non-path environments differ the most, and subsequently where troop foraging might show the greatest differences. This experimental setup is intended to represent a baboon group starting from a fixed location, such as a sleeping site. Adding a high-density path presents the group with a clearly advantageous foraging trajectory. We measure each individual’s intake of food over the simulation, as well as the distance from the high-density path to the center of the group.

We then set up a set of second foraging trials, where we fixed the group size and social structure and varied environmental structure. Two groups of 50 agents, one with a core of 45 agents and a second with a core of 5 agents, were made to forage on (i) landscapes in which the width and amount of food on the path were varied (Fig 2a), and (ii) landscapes in which the number and length of the paths were varied (Fig. 2b). For (i) we used a uniform random distribution to specify landscape structure, with path width varying between 10 and 100m, and amount of food in the path varying between 0.001 and 0.004 patches/m^2^. For (ii) we again used a uniform random distribution to specify landscape structure, varying the number of paths from 1 to 8, where the length of the paths where made smaller as the number of paths went up (e.g., the landscape with 2 paths had 2 paths each 1/2 the size of the one path landscape, 4 paths each 1/4^th^ the size of the one path landscape, … etc). For both sets of trials that varied an aspect of environmental structure, we generated 500 foraging landscapes and simulated foraging for small and large core groups, resulting in 1000 runs for each trial.

**Figure 2:**
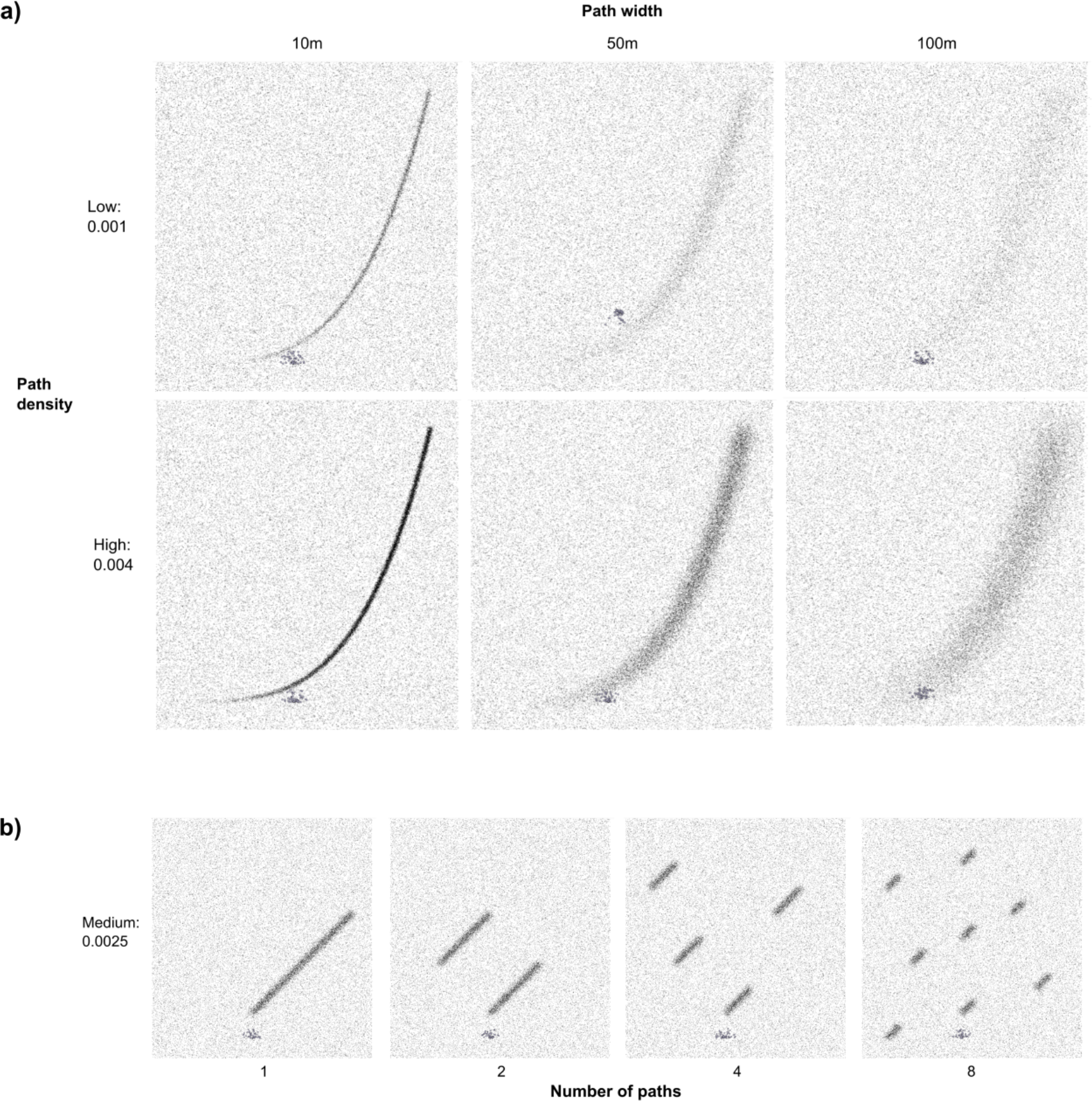
Range of resource landscapes used in the foraging trails. The high-density paths were added to a background of randomly distributed food patches by a) varying the width and density of food patches within a preset path, and b) varying the number and size of the paths in the landscape. The simulated group started each foraging trail located at the bottom middle of the landscape.

## Results

### Foraging efficiency: which group structures do better and under what conditions?

In a uniform habitat, groups with larger cores outperformed those with small cores, showing consistently higher food intake across the entire range of group sizes (Fig. 3a). When foraging in a landscape with a high-density path, however, we found that groups with smaller cores could sometimes outperform groups those larger cores across the range of group sizes, although they could also do much worse (Fig. 3b). Overall, foraging efficiency was higher under conditions in which a high-density path was present (Fig. 3).

**Figure 3:**
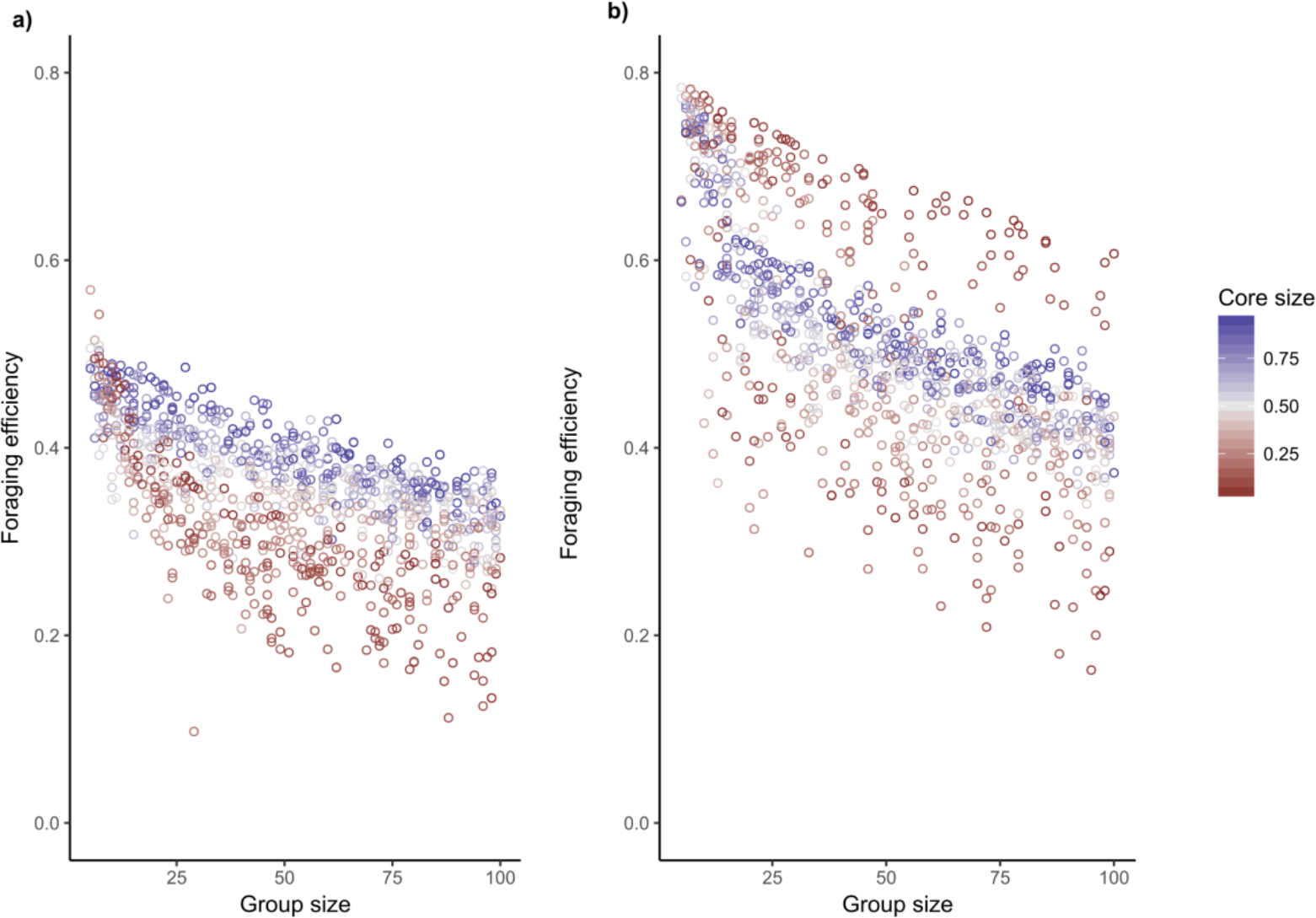
Rate of foraging intake under alternative influence structures, group size and landscape settings: a) uniform landscape, and b) high-density path on the landscape. Foraging intake is the percentage of total possible intake. The color of the points represents the percentage of the group that is part of the core, the remaining individuals are assigned to the periphery and follow a randomly specified core member.

As a further check on this, we compared the difference in foraging efficiency of groups of a given size and composition in the structured versus unstructured environment. This revealed that almost all combinations of group size and structure performed better in the environment with the high-density path. Nevertheless, groups with smaller cores apparently were able to benefit more from the presence of a high-density path than large core groups, and the strength of this effect increased with group sizes above 25 producing a bifurcation (Fig. 4).

**Figure 4:**
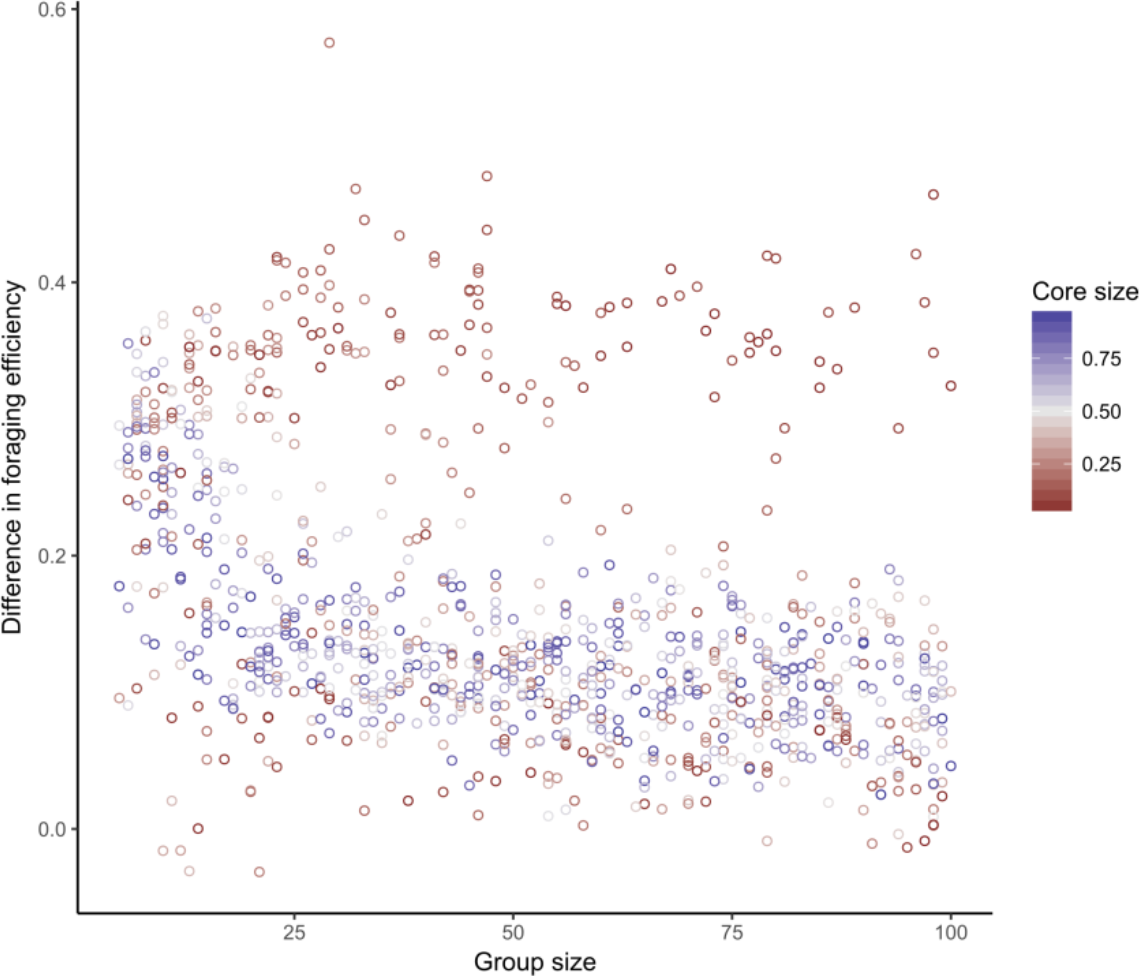
Difference in foraging efficiency across environments with and without the high-density path for a given group size and core-periphery structure.

This bifurcation can be explained by examining the groups’ distance from the high-density path Fig 5a). Small core groups that showed large positive differences in foraging efficiency (the upper part of the bifurcation) were also the ones that maintained close proximity to the high-density path (Fig. 5a). Although groups with large cores maintained looser proximity to the high-density path, groups of all sizes consistently remained within 200m of it. Larger groups with small cores often wandered very far from the high-density path resulting in reduced efficiency (Fig. 5a).

**Figure 5:**
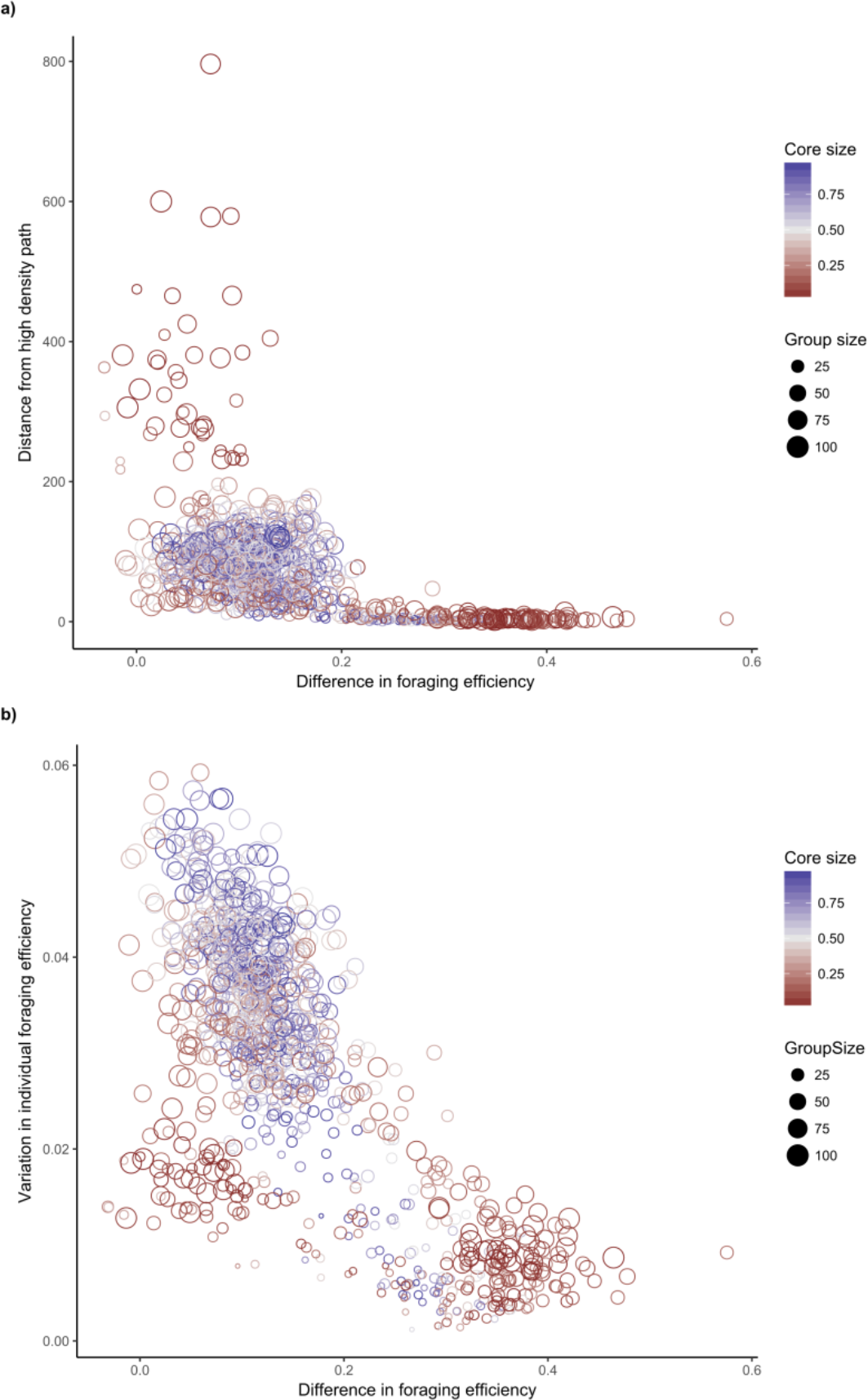
Resulting foraging patterns when group size and social structure are varied. The ability of groups to (a) maintain close proximity to the high-density path, and (b) the level of individual variation in foraging efficiency are compared to the ability of groups to take advantage of resource structure in the environment. The ability of groups to take advantage of resource structure is measured as the difference in foraging efficiency for each group between the high-density path and non-path environments (i.e., control). Point size represents the size of the group, and the color represents the size of the core within the group.

### Variability in foraging efficiency: do peripheral individuals benefit?

When we compared variability in individual level foraging efficiency, we found that large groups with large cores showed the highest intra-group variability in performance (Fig. 5b). As group size decreased, groups with large cores tended to show reduced individual variability along with increased foraging efficiency in the structured environment. For groups with small cores, there were two outcomes, that did not seem depend on group size (Fig. 5b). One outcome corresponded to small core groups that performed much better in the structured (high-density path) environment, while the other corresponded to small core groups that performed only marginally better in the structured environment. In both cases, there was lower intragroup variability compared to groups with large cores

### Varying environmental structures: what can groups with different structures exploit?

We then investigated how groups of a fixed size but different core-periphery structures responded to variation in environmental structure. We found that groups with small cores responded to both the size of the high-density path and amount of food it contained (Fig. 6ab, Table 1), whereas groups with large cores largely responded only to the amount of food (Fig. 6cd, Table 1). When overall foraging efficiency was compared, we found that groups with large cores tended to do better under most conditions (Fig. 6ef).

**Figure 6:**
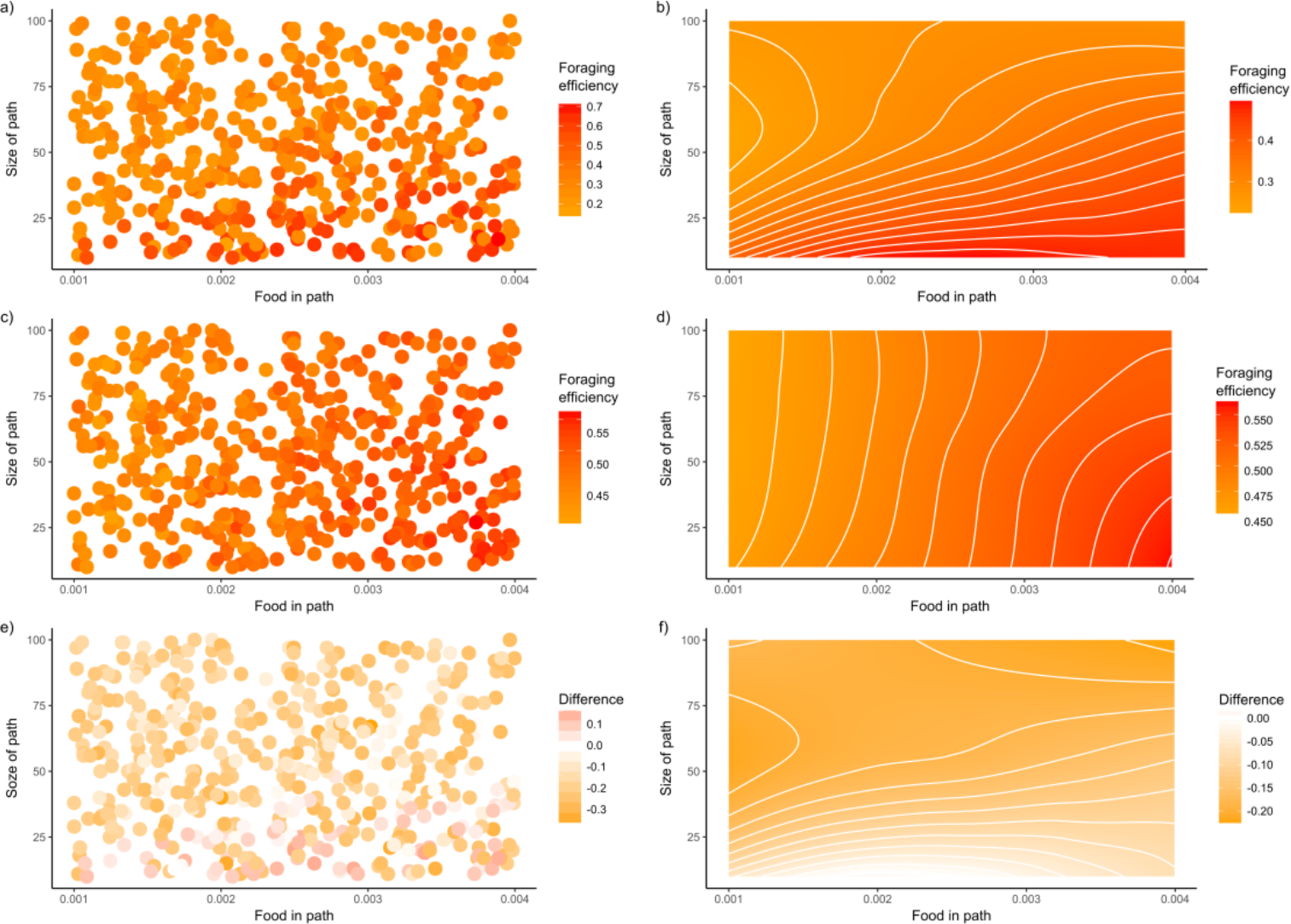
Foraging outcomes in resource environments with varying structure. Environmental structure was varied by adding a high-density resource path and altering the width and amount of food in the path. Two groups were made to forage in each resource environment: (a) a group of 50 individuals with 5 core members, with (b) modeled trend (loess), and (c) a group of 50 individuals with 45 core members, with (d) modeled trend (loess). The difference between the two groups under the same environmental conditions (e) and modeled trend (f) are presented.

**Table 1:** Linear model comparing the effect size of the amount and size of high density paths on foraging outcomes.

| Group | Standardized estimate (sd) |
| --- | --- |
| Large core group |  |
| Size of path | -0.15 (0.03) |
| Food in path | 0.76 (0.03) |
| Adj R<sup>2</sup> | 0.60 |
| Small core group |  |
| Size of path | -0.41 (0.04) |
| Food in path | 0.27 (0.04) |
| Adj R<sup>2</sup> | 0.25 |

When we varied the number and size of high density paths, creating a gradient from one long structure to many small structures, we found that groups with small cores had the ability to outperform groups with large cores only when there were a few large structures in the environment (Fig. 7, Fig. 2b). Otherwise groups with large cores consistently outperformed those with small cores.

**Figure 7:**
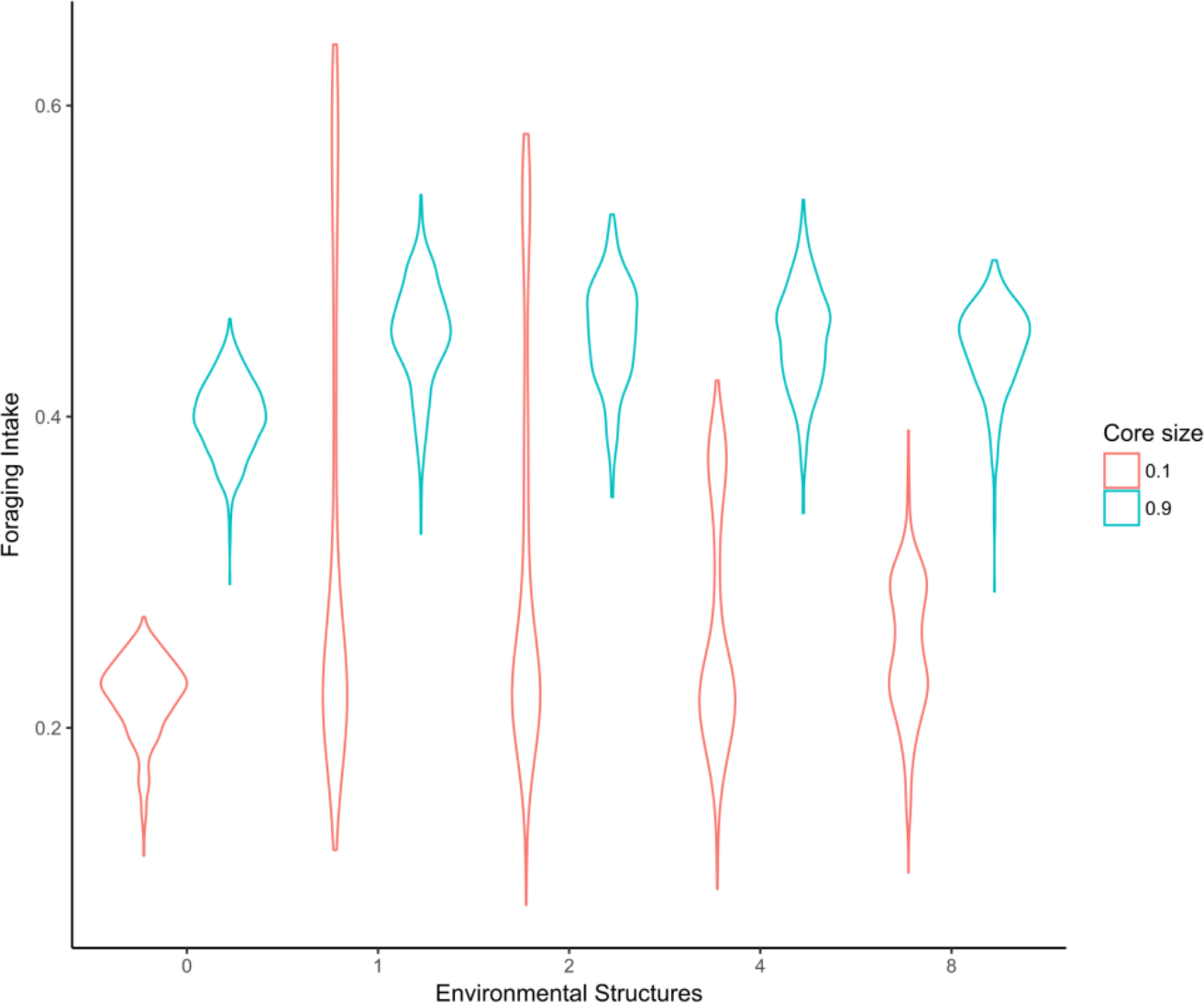
Foraging intake for large core and small core groups with varying the environmental structure. Environmental structure varied from no additional structure (0 = uniform distribution), to the addition of one long high-density resource path, to an increasing number of smaller and more numerous high density paths (e.g., 8 = 8 paths 1/8^th^ the size of the initial path). The distribution of foraging outcomes are represented by violin plots, where the width indicates the density of outcomes at a particular value of foraging intake.

## Discussion

Our results show that the structure of the resource environment can have a large impact on the functional outcomes of social influence structures, and accounting for environmental structure is thus an important consideration when attempting to understand the drivers of social influence patterns within baboon groups. More specifically, our simulations make the prediction that the development of homogenous influence structures (i.e., decentralized groups with large cores) will be favored in homogenous resource environments. For more structured resource environments, however, our simulations suggest something more nuanced as the outcomes are likely to depend on both the degree to which centralized structures hurt the group when it fails to locate resources in the environment (i.e., the costs of reduced detection), and the exact nature of environmental structure. When our simulated baboon group was presented with a generally homogenous environment with a single structured component (i.e., our high-density path), the failure to detect the path, as a consequence of possessing a small core of influential animals, incurred a high cost (Fig. 3a). When groups with small cores were presented with a more heavily structured landscape (i.e., several small high-density paths), the costs of missing one structural component (i.e., the difference between high performing and low performing small core groups) was reduced (Fig. 7). In the case of groups with larger cores, foraging benefits remained similar across all resource structures. Path width also interacted with group structure: in landscapes where the path width of the resource was relatively narrow, the added persistence of small core groups in maintaining proximity to such structures allowed such groups to forage more efficiently (Fig. 5a). Thus, small cores may be most effective under conditions when habitats are heterogeneous, with a few areas of high-density resources that are heavily restricted spatially.

Interestingly, and contrary to our original intuitions, we found that groups with smaller cores displayed lower variation in individual foraging intakes compared to groups with large cores, and this occurred regardless of whether groups with small cores detected the high-density path. More specifically, when groups with small cores found the high-density path, this resulted in both increased group-level foraging intake and decreased individual variability, suggesting that peripheral individuals benefited from the group’s closer proximity to the high-density path. When groups with small cores failed to find the high-density path, group-level foraging intake dropped, accompanied by a slight increase in individual variation, although this remained lower than for groups with large cores. One possible explanation here is that this reflects variation in travel speed: groups with smaller cores move faster across the landscape than those with larger cores, as the latter have a greater tendency to meander. As a result, peripheral individuals in groups with smaller cores may encounter new food sources more rapidly than peripheral individuals in slower, more meandering groups, and hence ensure inter-individual variation in foraging intake remains relatively low. For groups with large cores, we found that variation in individual foraging intakes decreased with decreasing group sizes, and was accompanied by an increase in group foraging intake. Overall, this suggests that smaller groups and lower inter-individual variation in foraging intake are both associated with shorter distances from the high-density path. This, in turn, suggests that smaller groups with larger cores are better able to take advantage of this form of highly concentrated environmental structure (Fig. 5ab).

More generally, our results conform to predictions that more centralized social groups, with influence structures tied to very few individuals, produce more extreme outcomes (Conradt and Roper, 2005). That is, groups with small cores either find and exploit the structure of the environment highly effectively, or they miss the high-density path completely and so fail to exploit it at all. Groups with larger cores, on the other hand, are highly effective at finding these kinds of environmental structure but are not as effective at exploiting it when they do so. Thus, variation in core size can be seen as a trade-off between the benefits of exploitation versus exploration (Fig. 5a).

In our simple model, there are no other mechanisms by which groups with smaller cores can increase their ability to detect environmental structure, nor for groups with larger cores to increase their effectiveness at exploiting of environmental structure (i.e., they have no means of maintaining tighter proximity to the path). As such, we have presented a form of null model, where our predictions are based solely on individuals that are foraging for local resources with a social bias in movement. Empirical data that deviates from these predictions can therefore help identify novel mechanisms by which baboon groups (and indeed groups of other species) increase their ability to detect and/or exploit environmental structure, and this in turn may be dependent on whether they possess a centralized (small core) or decentralized (large core) influence structure. Similarly, observational studies might also point to alternative social influence structures that have enhanced functional outcomes, i.e., those not neatly categorized as centralized or decentralized. Longitudinal studies might be most useful here, as the development of particular social influence structures could then be observed and enable the quantification of the relationship between environmental contexts (e.g., seasons) and social structures (e.g., movement bias) in baboon troops. Cross-sectional studies could also highlight differences between different groups within a population under differing environmental conditions, as well as cross-species comparisons (Reyna-Hurtado et al., 2017).

The *Papio* baboons offer great potential in this respect, as they are found throughout many differing environments, their evolutionary history is extremely well studied, and the different allotaxa show a variety of social structures that lend themselves to empirical tests of the kind suggested here (Barton et al., 1996; Henzi et al., 2009; Jolly, 2001; Patzelt et al., 2011; Schreier and Swedell, 2009; Snyder-Mackler et al., 2012; Weyher et al., 2014). Building on our simulated results, we may therefore be able to acquire an even better grasp on how social structure enables baboons to make the best use of space, time and energy.

## Acknowledgments

Thanks to Dietmar Zinner and Julia Fischer for the invitation to present at the Gottingen symposium on the future of baboon studies on which this paper is based, and to Cliff Jolly and Jeff Rogers for discussions of issues relating to all things baboon.

## Supplementary material

### Simulation Model

Full model code is available from: github.com/tbonne/Functional-Influence-Structures

### Clustering analysis

To aid in the interpretation of the simulated foraging outcomes a clustering approach was used. Foraging outcomes for each simulated group was measured by: 1) the difference in foraging efficiency of this group when foraging on landscapes with and without a high density path, 2) variation in individual foraging within the group, and 3) the distance between the high density path and the group. The function NbClust (Charrad et al., 2014) was used to determine the optimal number of clusters, using canberra distance, the Ward D2 method (Murtagh and Legendre, 2014), and 26 indices to test the validity of the choice of clustering (choosing the number of clusters selected by the majority of the incises). The optimal number of clusters in foraging outcomes was 4, and can be visualized in figure S1-ab. The degree to which these outcomes correspond to group size and core size within the groups can be seen in figure S1 c.

**Figure S1:**
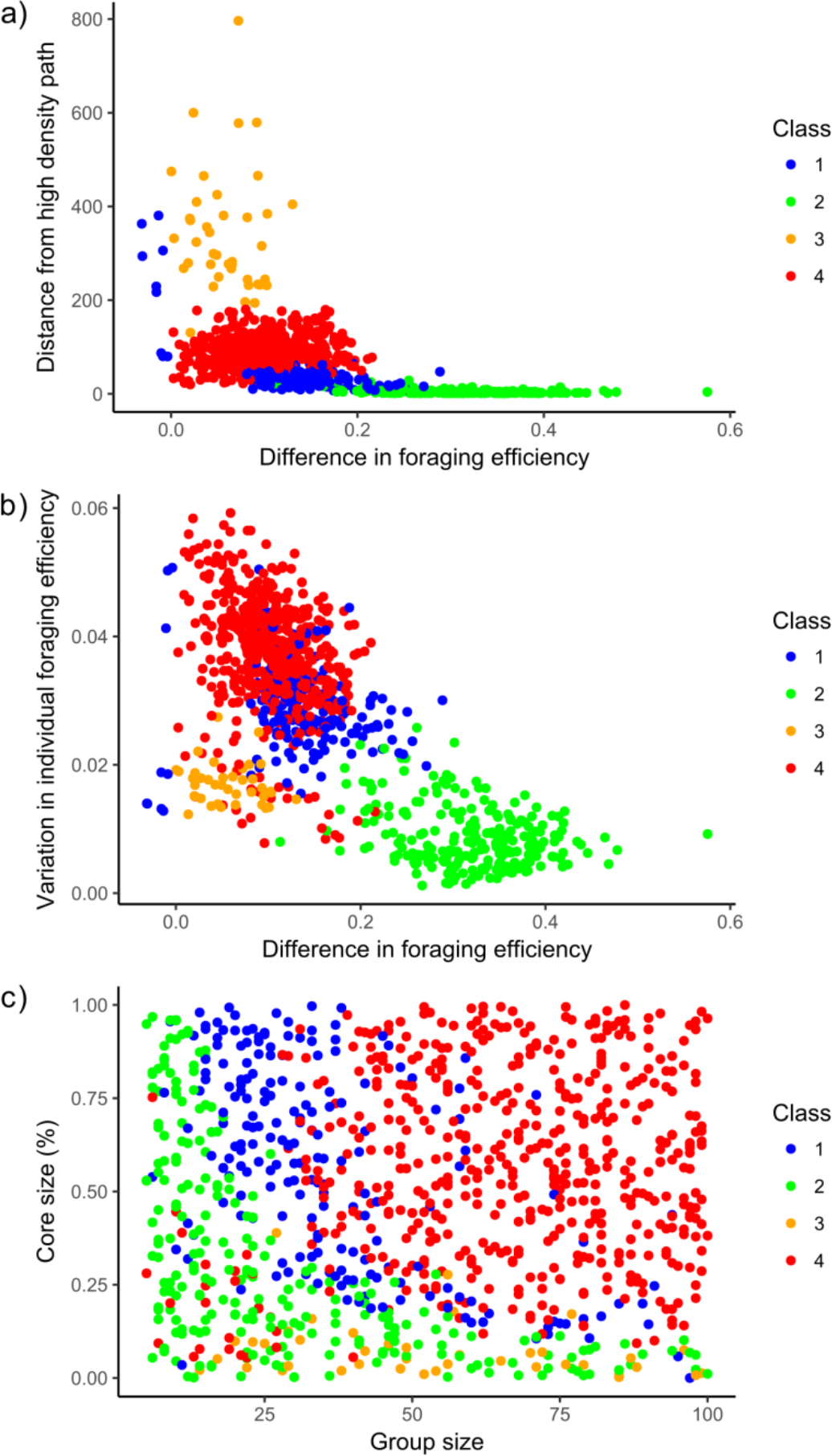
Clustering of foraging outcomes: a-b) identified classification of outcomes based on foraging efficiency, variation within group foraging, and distance maintained from the high density resource path. Plot c) displays how the categories of outcomes compare with the group size and core size measures.

